# RNA-seq 2G: online analysis of differential gene expression with comprehensive options of statistical methods

**DOI:** 10.1101/122747

**Authors:** Zhe Zhang, Yuanchao Zhang, Perry Evans, Asif Chinwalla, Deanne Taylor

**Affiliations:** Department of Biomedical and Health Informatics, The Children’s Hospital of Philadelphia, Philadelphia, Pennsylvania, 19104, USA

## Abstract

RNA-seq has become the most prevalent technology for measuring genome-wide gene expression, but the best practices for processing and analysing RNA-seq data are still an open question. Many statistical methods have been developed to identify genes differentially expressed between sample groups from RNA-seq data. These methods differ by their data distribution assumptions, choice of statistical test, and computational resource requirements. Over 25 methods of differential expression detection were validated and made available through a user-friendly web portal, RNA-seq 2G. All methods are suitable for analysing differential gene expression between two groups of samples. They commonly use a read count matrix derived from RNA-seq data as input and statistically compare groups for each gene. The web portal uses a Shiny app front-end and is hosted by a cloud-based server provided by Amazon Web Service. The comparison of methods showed that the data distribution assumption is the major determinant of differences between methods. Most methods are more likely to find that longer genes are differentially expressed, which substantially impacts downstream gene set-level analysis. Combining results from multiple methods can potentially diminish this bias. RNA-seq 2G makes the analysis of RNA-seq data more accessible and efficient, and is freely available at http://rnaseq2g.awsomics.org.

## INTRODUCTION

RNA-seq is by far the most advanced technology for investigating gene expression (1). It uses massively parallel sequencing to obtain millions of short reads from each cDNA library (2). After the sequencing reads are aligned to a reference genome and transcriptome, the number of reads mapped to each gene is used as a digital measurement of gene expression. Applications of RNA-seq data include, but are not limited to, alternative splicing, gene fusion, and allele-specific expression (3). However, its most common application is the detection of genes that show differential expression (DE) between sample groups.

Many statistical methods have been applied to DE analysis of RNA-seq data. Some were developed specifically for RNA-seq data, such as edgeR (4) and DESeq (5), and others were adapted from methods previously used for microarray data, such as limma (6) and SAMSeq (7). Methods using the original matrix of gene-level read counts as input often assume that counts have negative binomial distribution across samples while methods using log-transformed data are most commonly based on a normal distribution. Other methods assume Poisson-like, nonparametric, or multivariate distribution. Finally, metaseqR (8) uses a meta-analysis-like process to combine results of multiple methods. Numerous studies have compared different DE methods in terms of their technical similarities and precision of detecting DE genes (9). However, no detailed guidelines are available for data analysts to objectively select an optimal method for a specific data set analysed for a specific purpose.

Two follow-up strategies are commonly utilized once DE genes are identified from an RNA-seq data set. The first is to pick a small number of top DE genes and experimentally validate them using methods like quantitative PCR. The validated DE genes can be further investigated as biomarkers, disease classifiers, or regulators of downstream events. The second strategy is to select a larger number of DE genes and run an analysis to relate these genes to known biological processes, which is often called gene set enrichment or over-representation analysis. If the method selected for DE analysis has a systemic bias towards certain types of genes, the bias can be amplified by the gene set-level analysis. An example of such bias is that on average, more sequencing reads will be mapped to longer genes, which gives them a higher statistical power to be detected as DE genes.

We developed a user-friendly web portal, RNA-seq 2G, for users to choose any of over 25 DE methods and simultaneously run them online. The results of multiple methods can be visualized, compared, and combined online, or downloaded for other follow-up analyses. All DE methods were re-implemented as R functions to support the offline analysis of large or confidential data sets. Expression data from human tissues from the GTEx project (10) was used in this report to demonstrate our preliminary findings from comparing these methods.

## MATERIAL AND METHODS

### Evaluation and implementation of DE methods

We evaluated and tested over 40 DE methods (4–8, 11–25). About one-third of them were not selected for various reasons. For example, RBM is essentially the same as limma, ShrinkSeq is impractically slow, and DEGseq is over-sensitive. The remaining two-thirds of the methods were reimplemented in the DEGandMore package written with R, so all of them have the same input and output formats (http://github.com/zhezhangsh/DEGandMore).

These DE methods were also made available through the RNA-seq 2G web portal. It takes seconds to hours for the web portal to finish running the DE methods on a typical RNA-seq data set (10–20 samples). Correspondingly, we grouped the methods into four speed groups. About half of them are in the “Fast” group, where each takes less than 30 seconds to finish. The “Medium” methods usually take 0.5 to 5 minutes, and the “Slow” methods take up to 30 minutes. The “Slower” methods require even more time, so users will need to submit an analysis and retrieve its results later through an analysis ID. Other attributes of the DE methods are listed in Table 1. For example, some methods use the original read counts as input while others use log-transformed data. Additionally, about one-third of the methods cannot perform paired tests and some methods do not have their own normalization procedure, making them reliant on the normalization options available via RNA-seq 2G. Another main difference between the methods is their data distribution assumption, which is most commonly a normal or negative binomial distribution.

**Table 1.** DE methods available through RNA-seq 2G web portal and their attributes. (*Method*: method name; *Speed*: how fast is the method; *Paired*: whether the method supports paired test; *Logged*: whether the method uses read count and log-transformed data as input; *Norm:* whether the method has its own normalization procedure; *Distribution*: the data distribution assumed by the method; *Test*: the statistical test used by the method; and *R Function*: the core R function {package} of the method).

| Method | Speed | Paired | Logged | Norm | Distribution | Test | R Function |
| --- | --- | --- | --- | --- | --- | --- | --- |
| **StudentsT** | Fast | Yes | Yes | No | Normal | Student's t test, equal variance | TDist {stats} |
| **limma** | Fast | Yes | Yes | No | Normal | Empirical Bayes moderation | ebayes {limma} |
| **PoissonSeq** | Fast | Yes | No | Yes | Poisson-like | Poisson goodness-of-fit | PS.Main {PoissonSeq} |
| **edgeR** | Fast | Yes | No | Yes | Negative binomial | Exact/Likelihood ratio | exactTest {edgeR} |
| **DESeq2** | Fast | Yes | No | Yes | Negative binomial | Generalized linear model | DESeq {DESeq2} |
| **ABSSeq** | Fast | Yes | No | Yes | Negative binomial | Sum of counts | callDEs {ABSSeq} |
| **Ballgown** | Fast | Yes | No | Yes | Normal | F-test | stattest {ballgown} |
| **BGMix** | Fast | Yes | Yes | No | Normal | Bayesian mixture model | BGMix {BGMix} |
| **PLGEM** | Fast | Yes | No | No | Normal | Power Law Global Error Model | run.plgem {plgem} |
| **SAM** | Fast | Yes | Yes | No | Normal | Alternative t test with permutation | samr {samr} |
| **voom** | Fast | Yes | No | Yes | Normal | Empirical Bayes moderation | voom {limma} |
| **WelchST** | Fast | Yes | Yes | No | Normal | Welch's t test, unequal variance | TDist {stats} |
| **EBSeq** | Medium | No | No | Yes | Negative binomial | Empirical Bayesian | EBTest {EBSeq} |
| **LMGene** | Medium | Yes | No | No | Normal | Linear model & glog transformation | genediff {LMGene} |
| **NOISeq** | Medium | No | No | Yes | Nonparametric | Empirical Bayes | noiseqbio {NOISeq} |
| **RankProd** | Medium | Yes | Yes | No | Nonparametric | Rank product | RP {RankProd} |
| **SAMSe** | Medium | Yes | No | No | Nonparametric | Wilcoxon with resampling | SAMseq {samr} |
| **Wilcoxon** | Medium | Yes | Yes | No | Nonparametric | Wilcoxon signed-rank test | wilcox.test {stats} |
| **ALDEx2** | Slow | Yes | No | Yes | Multivariate | Welch's t, etc. | aldex {ALDEx2} |
| **baySeq** | Slow | Yes | No | Yes | Negative binomial | Empirical Bayesian | getLikelihoods {baySeq} |
| **DEXUS** | Slow | No | No | Yes | Negative binomial | Bayesian framework by an EM algorithm | dexus {dexus} |
| **NBPSeq** | Slow | No | No | Yes | Negative binomial | Robinson/Smyth exact NB test | nbp.test {NBPSeq} |
| **sSeq** | Slow | No | No | No | Negative binomial | Shrinkage Approach of Dispersion Estimation | nbTestSH {sSeq} |
| **BADER** | Slower | No | No | Yes | Poisson-like | Bayesian | BADER {BADER} |
| **metaseqR** | Slower | No | No | Yes | Meta-analysis | Simes method of combining p values | combine.simes {metaseqR} |
| **tweeDEseq** | Slower | No | No | Yes | Poisson-like | Poisson-like | tweeDE {tweeDEseq} |

### System architecture of RNA-seq 2G

RNA-seq 2G has two major elements. The first is the DEGandMore R package, and the second is a Shiny app hosted by a Shiny web server running on an AWS (Amazon Web Service) instance. This framework provides three options to run DE analysis. The easiest way is to submit an analysis to the web portal through a web browser (http://rnaseq2g.awsomics.org). Alternatively, R users can install the DEGandMore package and run the analyses locally. The local results can later be uploaded to the web portal for follow-up analyses. To keep their data confidential, users can initiate and run a new AWS instance based on its AMI (Amazon Machine Image).

### Using the RNA-seq 2G web portal

Each time a user visits the web portal via a web browser, a random analysis ID is generated. To submit an analysis, the user needs to upload an integer matrix of read counts with unique gene identifiers as row names and unique sample identifiers as column names. The matrix can be uploaded as an R object, Excel table, or tab-delimited text file. The second step is to group the samples for DE analysis and specify parameters such as the normalization method and whether to run a paired test. The last step is to select the DE method(s) to use and then submit the analysis to the server.

The results of finished analysis will be available immediately and saved on the server for later. The output of all DE methods is a six-column matrix with the same number of rows/genes. Columns 1 and 2 are group means of normalized read counts, column 3 is the mean difference, column 4 is the log2-ratio (fold change) of group means, column 5 is the statistical significance (p value) of the mean difference, and column 6 is the corresponding false discovery rate calculated by the Benjamini & Hochberg method (26). Each DE method has its own way of calculating the p values, based on data distribution, sample permutation, or posterior probability. Results of multiple DE methods can be interactively visualized, compared, and combined online.

Also available at the web portal are a detailed user manual and step-by-step instructions for submitting an analysis. Sample data and results that can be loaded for demonstration, or downloaded in various file formats. No user registration is required and an email address is optional and will only be used for notifying users of the completion of their analysis.

### Comparison of DE methods

DE methods were compared using a subset of the GTEx data (10), including samples of hippocampus and hypothalamus, two brain subdomains. The rand index of the two groups equals to 0.86, suggesting that they have somewhat similar transcriptomes. To reflect the typical sample size of RNA-seq data sets, six samples were randomly selected from each group for DE analysis. This was repeated 10 times to ensure that the results are consistent.

## RESULTS

### Gene-level analysis

The number of DE genes (p < 0.01) identified from the hippocampus vs. hypothalamus comparison by different methods ranged from 6 (BADER) to 2,789 (sSeq). Such a difference is likely derived from the statistical test used by DE methods and has little association with their assumption about data distribution. For example, DESeq2 identified 10 times more DE genes than baySeq, although both of them are based on a negative binomial distribution. We suggest that the relative ranking of p values is a more informative statistical index for comparing DE methods. For example, while the ranges of p values differed by over 2 orders between Student’s t test and SAM, their p value rankings were almost identical (Spearman’s correlation coefficient = 0.999) because SAM is simply a more conservative variation of t test.

Principal component analysis using p value rankings completely split DE methods based on their assumption about normal and negative binomial data distributions (Figure 1), although methods based on the same distribution still have plenty of disparity. For example, while both edgeR and DESeq are based on the negative binomial distribution, they had less than 85% overlap between their top-500 DE genes. Nonparametric methods have even less similarity to each other since they are not based on any data distribution and use different statistical tests, from rank sum (Wilcoxon) to rank product (RankProd).

We took a closer look of two genes with high-level expression in both hippocampus and hypothalamus (Figure 2). SLC6A11 encodes a neurotransmitter transporter. Its expression levels were perfectly split between the two groups and about 5-fold higher in hypothalamus on average. As a result, all DE methods ranked it as one of the top-100 DE genes. The other gene, PLP1, had higher average expression in hippocampus, but was not fully split between the groups. Consequently, different methods vastly disagreed on the significance of its DE, ranking it from top 1% (PLGEM) to bottom 1% (baySeq).

**Figure 1.**
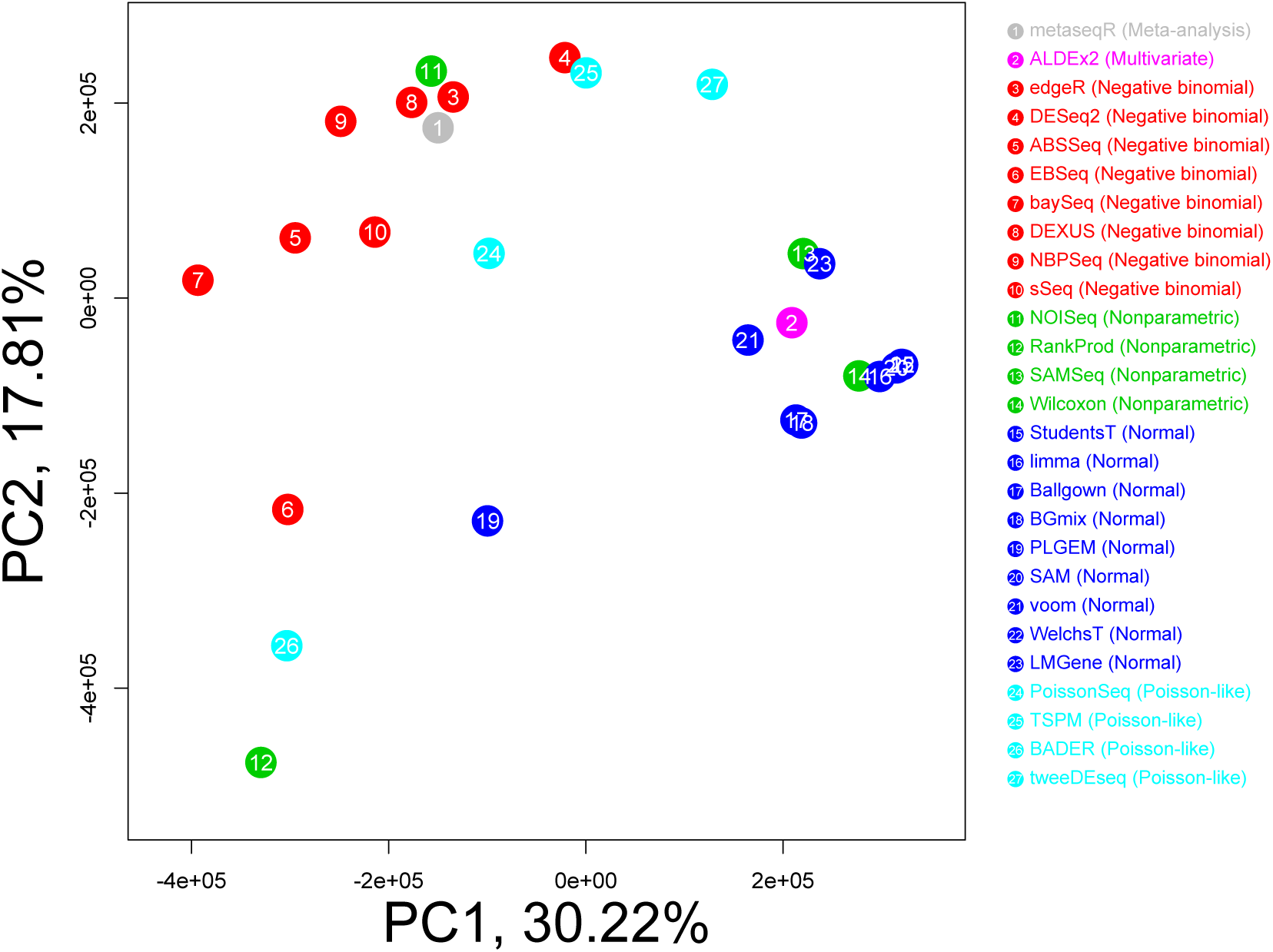
PCA (principal component analysis) was applied to the p value rankings of all genes to compare the global patterns of different methods. The methods are colored based on their data distribution assumptions.

**Figure 2.**
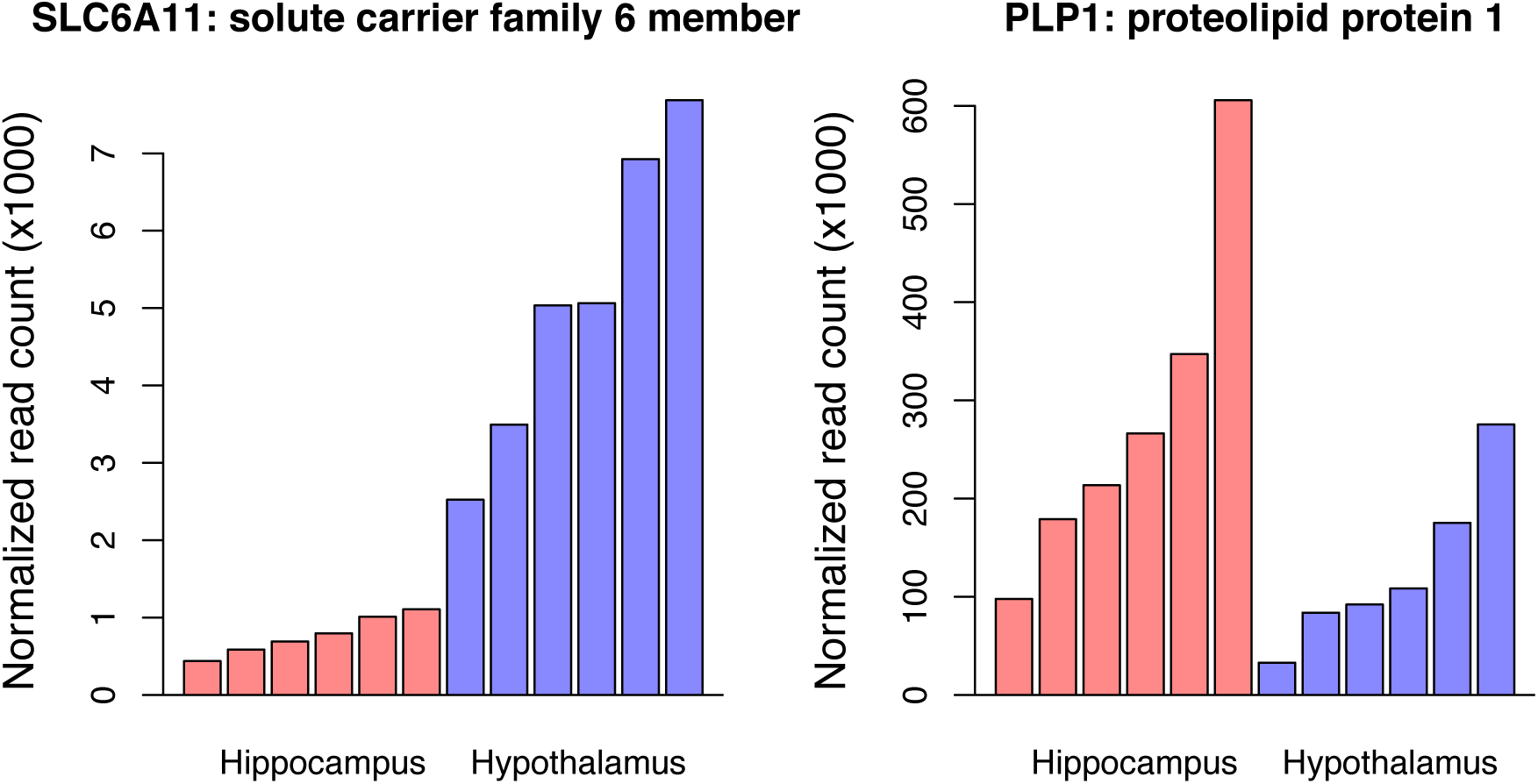
The normalized read counts of two example genes.

### Gene set-level analysis

We postulated that gene length is a source of bias during DE analysis. To perform a comparison of short vs. long genes, we selected two groups of protein coding genes from those having at least medium expression levels in both groups. The “short” group includes the 25% of genes having the shortest length (mean = 1.3kb), while the “long” group includes top 25% of genes having the longest length (mean = 5.7kb). On average, a long gene had a 50% higher read count than a short gene, but the short genes had higher expression than the long genes according to length-adjusted read counts. All DE methods except RankProd favoured longer genes. For example, both DESeq2 and edgeR were 20% more likely to give significant p values to a long gene than a short one.

At the gene set-level, the systemic bias of DE analysis is amplified. For example, genes of voltage-gated sodium channel activity (GO:0005248) have an average length over 5kb. It is indeed a favorite of gene set-level analysis according to our previous experiences. To run a simulation, we randomly selected genes from the “long” and “short” groups to form a collection of “pseudo” gene sets. Gene set enrichment analysis was applied to those gene sets and top DE genes ranked by each method. Gene sets of long genes were much more likely to obtain significant results than those of short genes according to most DE methods, such as tweeDEseq (162-folds), DESeq2 (17-fold), edgeR (3-fold), and limma (8-fold). On the other hand, RankProd (102-fold) and Wilcoxon (5-fold) favored gene sets of shorter genes. Methods showing no strong bias include NBPSeq (0.82-fold), PoissonSeq (1.06-fold) and metaseqR (1.08-fold).

## DISCUSSION

Using simulated or experimental data sets, previous studies have compared DE methods of RNA-seq data by focusing on their sensitivity and overlapping of DE genes (26–30). Their results have not conclusively pinpointed a method that out-performed the others under all circumstances. We believe that the best practice of analyzing RNA-seq will only originate from the collective experience of data analysts and scientists after they apply various methods to various data sets, and closely examine the results. RNA-seq 2G will substantially accelerate this process by providing an objective comparison platform.

To develop RNA-seq 2G, we evaluated ∼40 methods for two-group DE analysis. Over 25 of them were validated and made available through a user-friendly web portal. It provides at least twice as many choices of DE methods as any previous studies. The cloud-based system in combination with the Shiny server minimized the effort involved in system administration and user interface design. We believe it is a practical paradigm for developing more web-based applications.

The number of DE genes found from the same data set can differ by 2-3 orders when the same p value cutoff was used, reflecting the broad disparity of sensitivity between methods. Therefore, it is more meaningful to compare the methods in terms of their relative ranking of genes. Given that so many different DE methods are available, it is also suggestive that hypothesis testing of individual genes should be applied with caution, in consideration of the objectivity of method selection. The RNA-seq 2G web portal will help users make informed decision about which method to pick via visualizing and comparing global patterns of results from multiple methods.

Methods making the same assumption about data distribution tend to obtain more similar results (Figure 1). Increasing sample size will improve the similarity of methods, but will not make distinctive methods converge. Scrutinizing individual genes explained the intrinsic disparity between methods as they evaluate different combinations of effect size and consistence of group difference differently (Figure 2). Data analysts should then select DE methods accordingly, based on which type of DE genes is more biologically meaningful in the systems they are investigating.

Unlike previous studies, we evaluated the impact of DE methods on downstream analysis at the gene set-level. Gene length was confirmed as a source of systemic bias during DE analysis. The bias was further amplified during gene set analysis to a degree that many DE methods favored gene sets of long genes by over 10 times. Such a bias should be carefully dealt with during integrative analysis of multiple data sets. Interestingly, combining results from multiple methods by metaseqR seemingly reduced the bias. The validation of such a strategy requires further investigation. Baseline expression level, GC content, and sequence mapability of genes are potentially other sources of systemic bias. These factors are related and distinct, and should be evaluated both individually and collectively in future studies.

In summary, the RNA-seq 2G web portal has by far the most comprehensive collection of DE methods, and will make the analysis of RNA-seq data more accessible and efficient. Our future studies will further compare DE methods in terms of their performance on different cell types and sample sizes, as well as their robustness to background noise and outlier samples. The impact of systemic bias on gene set analysis and strategies to remove such bias will also be further evaluated.

